## Supplementary material for "Pre-stimulus alpha power modulates trial-by-trial variability in theta rhythmic multisensory entrainment strength and theta-induced memory effect": Figure S

**Supplementary figures**

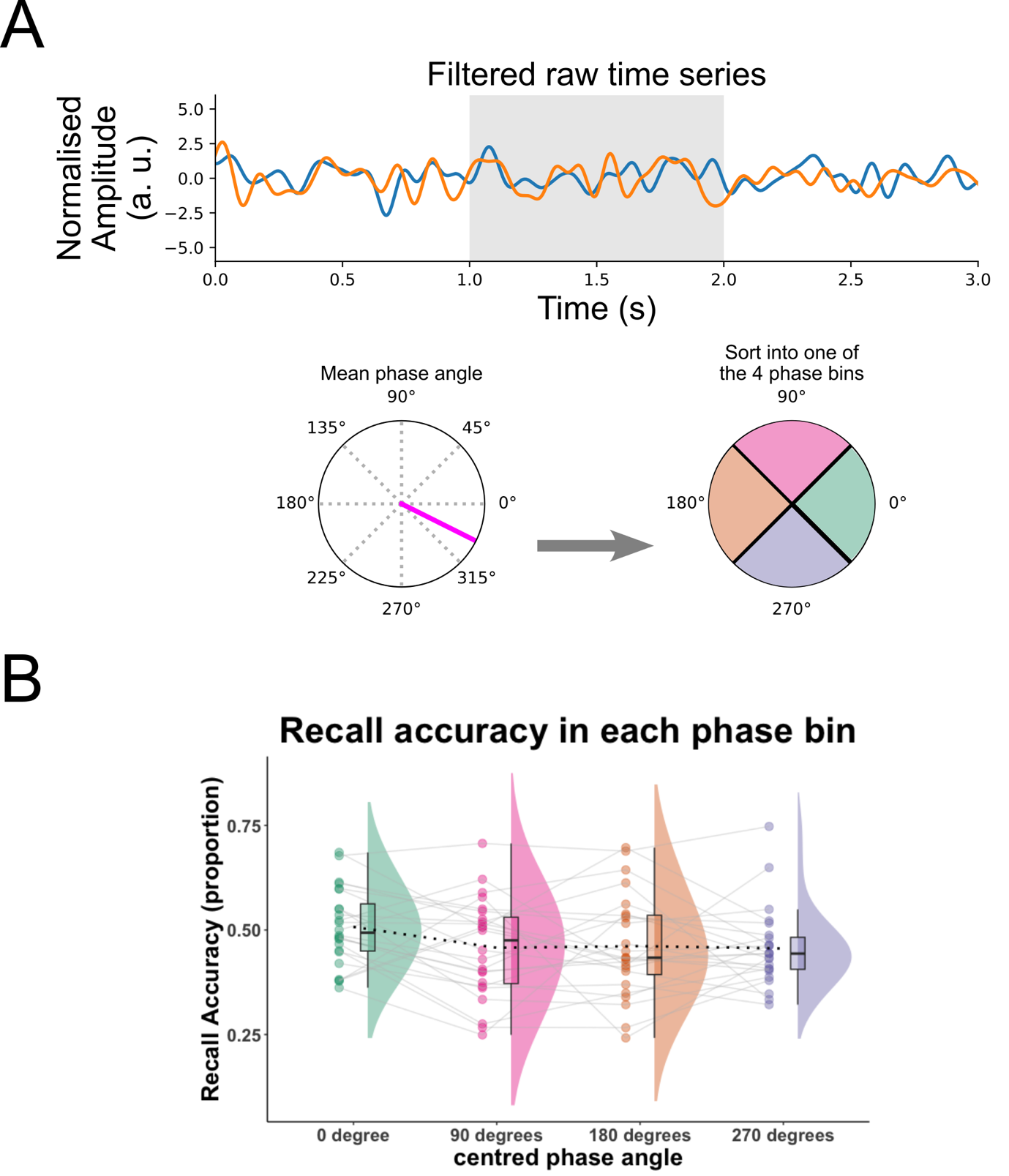

**Figure S1. Recall accuracy of each phase bin that consisted of trials which mean phase direction fell within one of the four phase bins that were centred at 0°, 90°, 180°, and 270°.** A, to replicate the previous findings (Wang et al., 2018), each trial was sorted into one of the four phase bins that were centred at 0°, 90°, 180°, and 270°, with a bin boundary of ±45º (bottom right), depending on the mean phase direction. B, the proportion of remembered trials was calculated for each phase bin the same way as was calculated in the two-bin analysis. A GLME was built with fixed factors ‘phase bin 1 vs 4’, ‘phase bin 2 vs 4’, ‘phase bin 3 vs 4’, and ‘subjective rating’, and random factors ‘subjects’, ‘movies’ and ‘sounds’, with control parameters of the optimx optimizer L-BFGS-B. The factor ‘phase bin 1 vs 4’ significantly predicted recall accuracy, *z* = 2.005, *p* = 0.045.

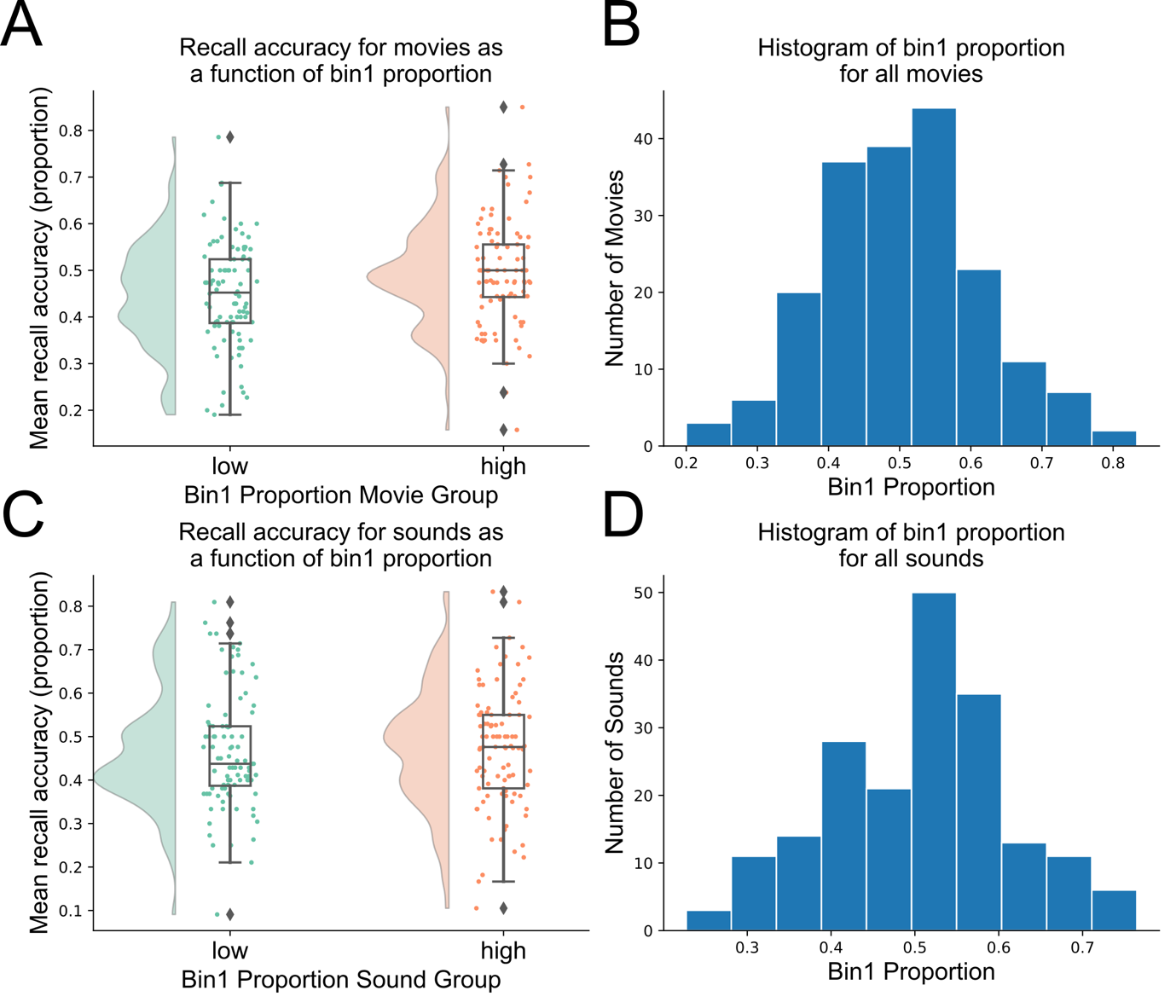

**Figure S2.** **The distribution of the proportion of movies or sounds that were assigned to phase bin 1 (centred at 0º, 2 bins).** A, recall accuracy was calculated for each movie by how many times the movie was successfully recalled among all 24 participants. Each movie was median split into either low bin 1 proportion group or high bin 1 proportion group. The mean recall accuracy was calculated for all the movies in each bin 1 proportion group. Movies in high bin 1 proportion group were remembered significantly more than the movies in the low bin 1 proportion group, *t*(190) = 2.649, *p* = 0.009. B, histogram of the distribution of the frequency of movies assigned to bin 1 among 24 participants. Bin 1 proportion in movies was normally distributed with mean bin 1 proportion 0.505 and median 0.5, suggesting most movies were assigned to bin 1 at chance level. C and D, same as A and B, but the bin 1 proportion and recall accuracy was calculated by how frequently a sound was assigned and remembered among 24 participants. No significant difference in mean recall accuracy between sounds in each bin 1 proportion group, *t*(190) = 0.378, *p* = 0.706. Bin 1 proportion in sounds was also normally distributed with mean 0.505 and median 0.5. Since better remembered movies were significantly more likely to be assigned to bin 2, movies were median split into better remembered and less remembered depending on the recall accuracy of movies among participants. Then we randomly sub-sampled equal trials from better remembered movies and less remembered movies for bin 1 and bin 2. The recall accuracy was still significantly better in bin 1 than in bin 2, *t*(23) = 3.082, *p* = 0.005.

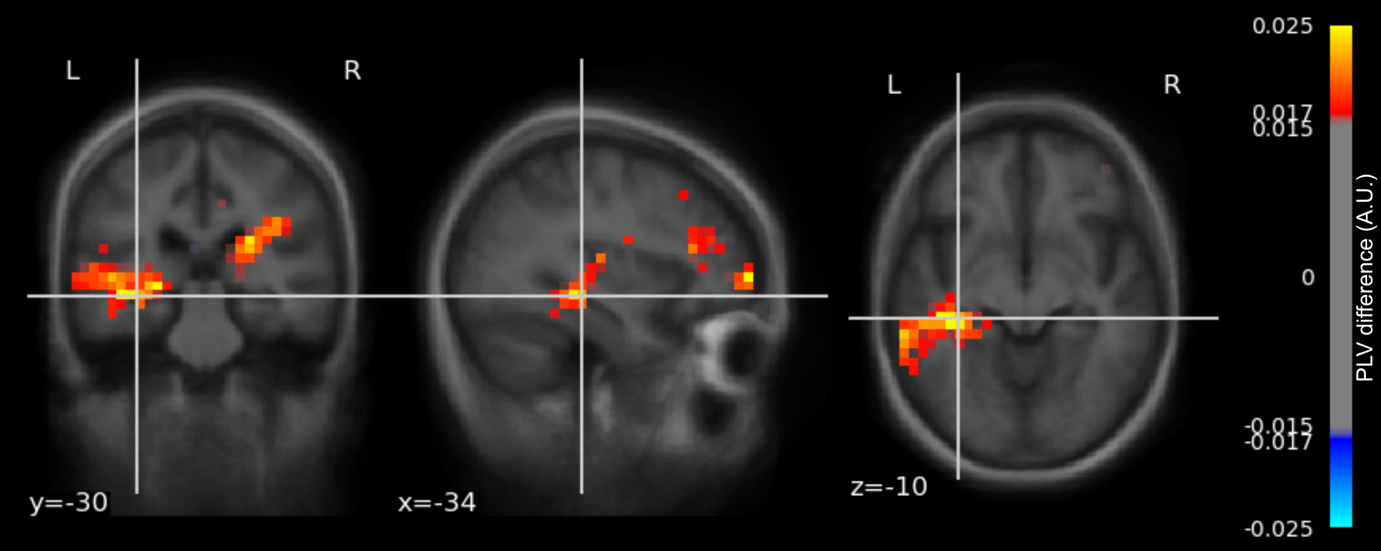

**Figure S3. Difference in pre-stimulus theta phase coupling between match and mismatch conditions.** Phase differences were calculated between the seed region (where the greatest pre-stimulus alpha power difference was found between match and mismatch conditions, MNI coordinates: -24, -75, 14) and the rest of the brain in theta frequency (3 – 8 Hz) for all data points across the whole epoch (-2 – 5 s). Phase locking value was calculated using the resultant vector length of the phase differences in the pre-stimulus time window -0.7 and -0.2 s, as well as the baseline time window 3.2 s and 3.7 s. The pre-stimulus PLV was normalised by the baseline PLV for each virtual sensor in each condition. The difference in the normalised PLV between the match and mismatch conditions was overlaid onto a standard MRI image in the MNI space. A cluster-based permutation one-sample t-test on the left hippocampus ROI showed that the PLV between the seed region and the left hippocampus increased significantly before the matched trials, as compared to the mismatched trials. The MNI coordinates highlighted here showed where the greatest t-value was in the permutation test: -34, -30, -10.
